# Hold out the genome: A roadmap to solving the cis-regulatory code

**DOI:** 10.1101/2023.04.20.537701

**Authors:** Carl G. de Boer, Jussi Taipale

## Abstract

Gene expression is regulated by transcription factors that work together to read cis-regulatory DNA sequences. The “cis-regulatory code” - the rules that cells use to determine when, where, and how much genes should be expressed - has proven to be exceedingly complex, but recent advances in the scale and resolution of functional genomics assays and Machine Learning have enabled significant progress towards deciphering this code. However, we will likely never solve the cis-regulatory code if we restrict ourselves to models trained only on genomic sequences; regions of homology can easily lead to overestimation of predictive performance, and there is insufficient sequence diversity in our genomes to learn all relevant parameters. Fortunately, randomly synthesized DNA sequences enable us to test a far larger sequence space than exists in our genomes in each experiment, and designed DNA sequences enable a targeted query of the sequence space to maximally improve the models. Since cells use the same biochemical principles to interpret DNA regardless of its source, models that are trained on these synthetic data can predict genomic activity, often better than genome-trained models. Here, we provide an outlook on the field, and propose a roadmap towards solving the cis-regulatory code by training models exclusively on non-genomic DNA sequences, and using genomic sequences solely for evaluating the resulting models.

## Background

In contrast to the protein code (**Box 1**), deciphering the cis-regulatory code remains one of the biggest unsolved problems in biology ^1–3^. Cis-regulation comprises the mechanisms by which DNA sequence determines transcription rates, and mRNA splicing and stability of nearby genes. Factors affecting mRNA splicing and stability has been reviewed recently elsewhere^4–7^. In this perspective, we will focus on how transcription factors (**TFs**) interpret cis-regulatory DNA sequences to control the rate of gene transcription. While the field has characterized many mechanisms used in cis-regulation (reviewed in ^8–12^), we lack sufficient understanding to combine this knowledge into predictive models^13^. At its core, cis-regulation is a quantitative process, relating DNA sequence to an amount of gene expression.

Accordingly, building a quantitative predictive understanding is vital to our efforts to design synthetic regulatory sequences (e.g. genetic circuits, cell-type specific gene therapies), and to understand how mutations to cis-regulatory sequences change their function. Perhaps nowhere is this more important than in regulatory sequence evolution, where we seek to ask how genetic variation impacts past, present, and future gene regulatory activity, and, consequently, organismal phenotype^14–16^. For instance, most of the common genetic variation associated with human traits (including diseases) appears to be regulatory, changing when, where, and the degree to which genes are expressed ^17,18^. Because of the astronomical number of potential cis-regulatory alleles, experimentally determining how they impact expression of nearby genes in every cell type is not feasible. However, a comprehensive cis-regulatory model would enable us to predict variant function, and could be equally applied to common, rare, *de novo*, and hypothetical variation, shedding significant light onto the mechanisms underlying diseases and other traits. A quantitative model will also be critical for fully understanding the molecular mechanisms of gene regulation, since it enables us to gauge the relative contributions of the different molecular mechanisms involved in driving transcription.

### Box 1

Contrasting the protein and cis-regulatory codes.

We have made great strides in deciphering the protein code, and while there are many parallels between the protein and cis-regulatory codes, there are also key differences. At the lowest level, proteins are encoded in the genome by the Genetic Code, which is shared amongst almost all life on Earth. The Genetic Code is essentially a discrete deterministic mapping between each triplet codon sequence and the amino acid to be incorporated into the protein. In contrast, the cis-regulatory code is quantitative, with TFs recognizing and binding to the DNA in proportion to their sequence affinities and cellular concentrations. Some regulatory proteins, like the histones that comprise nucleosomes, have extremely weak DNA sequence binding preferences that are nonetheless important to their function^19^.

At higher levels, proteins fold into a 3-D shape that determines their function. How a protein folds is relatively deterministic as well, with the same amino acid sequence folding into the same protein in every cell type of the body, and even across distantly related species. This enables repurposing of useful genes, like the Green Fluorescent Protein, across species. The conservation of folding biochemistry also enables us to use examples of structures of bacterial proteins to learn how human proteins fold. Indeed, the field has made significant progress in predicting protein structure from amino acid sequence recently, with ML programs like AlphaFold now able to predict protein structure from amino acid sequence accurately^20^. In contrast, higher order cis-regulatory features are much less deterministic and vary by species, cell type, cell state, and genomic locus because they depend on the concentrations and identities of the transcriptional regulators and the relative arrangement of the cis-regulatory elements along the DNA. This complexity makes predicting genome regulation from DNA sequence - let alone changes in regulation resulting from mutations - a major challenge. However, even for protein structures, predicting mutation effects remains a major challenge^21,22^.

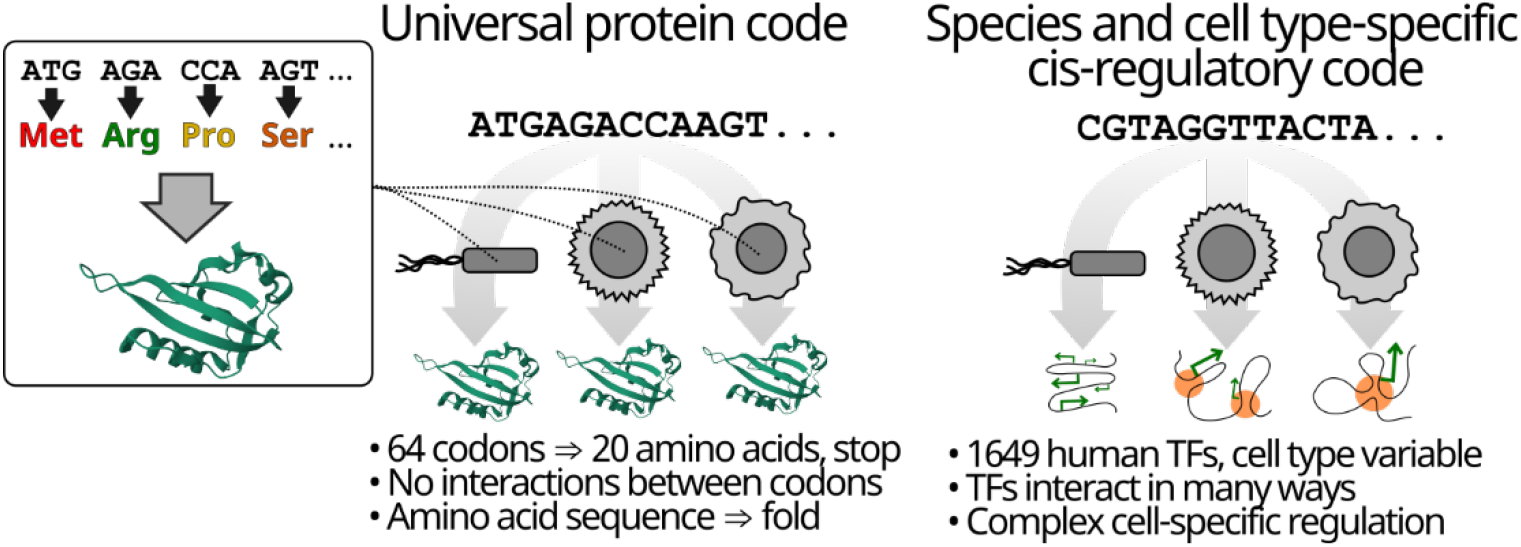

The multi-layered and convoluted nature of the cis-regulatory code partly explain why it remains unsolved. Cis-regulation is determined largely by TFs, proteins that recognize specific sequences in the DNA via their affinities to different DNA sequences (**Fig. 1A**). To a first approximation, cis-regulation appears to be quite simple, with the activities of most elements being consistent with a “billboard” model of gene regulation^23,24^, where the overall activity of a sequence can be reasonably approximated as the sum of its parts^25^. Recent studies have found that most regulatory sequences have 5-6 strong TF binding sites, with little preference for specific positioning or orientation between them^24^. This is consistent with the biochemistry of TFs, where TFs mutually compete with nucleosomes and their effector domains appear to be largely unstructured and interact transiently and in a variety of ways with co-regulator proteins ^26–28^ (**Fig. 1B**). Further, cis-regulation also appears to be hierarchical, with some TFs having outsized impact in any particular cell type^29,30^, but with many shared features between cell types as well^30,31^. While most gene expression can be explained with a billboard model, there are also clearly positional effects of TF binding as well that can modify TF activities ^25,32–37^, substantially complicating things. Finally, there is feedback between the chromatin state and transcriptional regulation^38^. Here, histone modifications and CpG dinucleotide methylation are added to chromatin by TFs and transcription^38–41^. These same modifications can then, in turn, modify how TFs read the DNA^42–44^. CpG methylation, and some repressive chromatin modifications such as H3K27 trimethylation can also persist through cell division^44–48^ to form a memory of past TF activity. Finally, adjacent regulatory elements interact via 3-D looping^49–51^, insulation^52^, competition^53–55^, and condensate formation^56,57^ to regulate the expression of nearby genes. There may also be hitherto undescribed mechanisms that further complicate cis-regulation. A predictive model of cis-regulation requires accurately capturing how all these layers of regulation work together.

**Figure 1:**
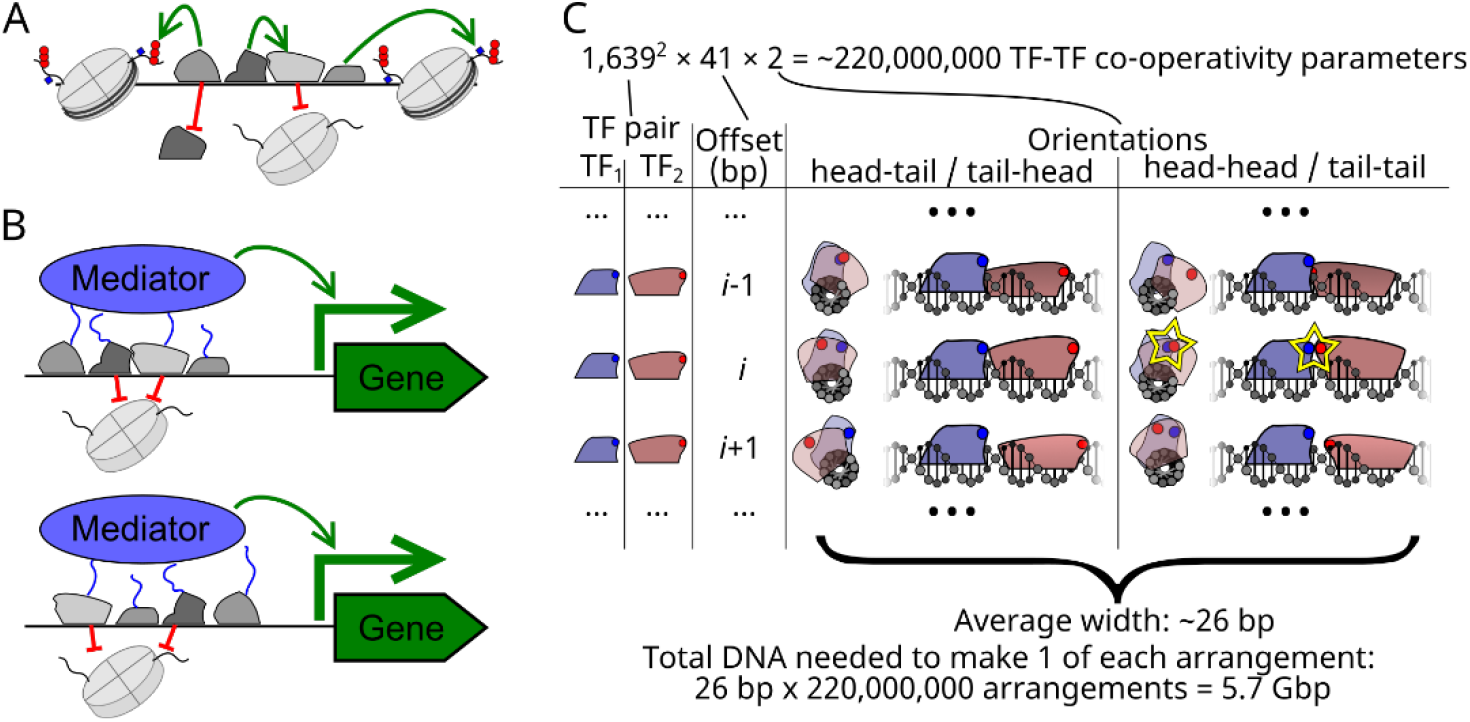
Cis-regulation requires learning hundreds of millions of parameters. (**A**) Some of the potential interactions between TFs (small amorphous blobs), DNA (black line), and nucleosomes (cylinders; histone modifications as small red circles). (**B**) At higher levels, regulatory information is compressed into few pathways via shared interactions with common factors (e.g. nucleosomes and Mediator). (**C**) TF-TF cooperativity includes ∼220 million parameters that impact TF binding. For each orientation, the TFs (coloured blobs) are shown bound to DNA as viewed as a cross-section of the DNA (left) and along the DNA’s length (right). A star highlights the cooperative interaction.

A second reason why cis-regulation remains unsolved is the very large number of parameters needed to describe the process. TFs can bind to DNA individually and in concert (**Fig. 1A**), interacting with each other both negatively, for instance by competing with each other for binding to overlapping DNA sequences, and positively, for instance by competing with nucleosomes^58^, and via biochemical cooperativity. Biochemical cooperativity is often mediated through direct physical interactions between TFs that are bound to adjacent DNA sequences that increase the affinity of one or both of the TFs to the DNA ^36,59^ (**Fig. 1C**). Because of the strict orientation, spacing, and DNA rotational requirements (e.g. each base pair results in a ∼34° rotation around the DNA double helix), such interactions often occur at only one of the many possible relative arrangements of the TF pair, although less stringent interactions are possible as well^59–63^. Since any of the ∼1,639 human TFs could potentially interact with any other, we estimate that there are approximately 220 million parameters needed to explain TF-TF cooperative interactions (**Fig. 1C**). However, this is likely to be an underestimate as this does not account for TF isoforms and post-translational modifications, and interactions with cofactors, which differ by cell type and are likely to modify these interactions. Systematic query of the TF-TF interaction space revealed that approximately 3% of TF pairs interact in specific binding site arrangements^37^, indicating that this parameter space, though sparse, is nonetheless important. Taken together, the multi-layered nature of the regulatory code, and the vast number of parameters required to define it makes creating comprehensive cis-regulatory models a major challenge.

There have been several major successes in creating genome-trained models that predict cis-regulatory activity from DNA sequence alone. Many of the recent advances rely on deep neural networks ^60,64–70^, which are sufficiently complex to capture many of the parameters of cis-regulation. For instance, the BP-NET model trained on ChIP-nexus data learned an interaction between Nanog and other TFs that followed a periodic dependency consistent with the helical period of DNA^60^. The Enformer model, which was trained to predict thousands of chromatin profiles across 200 kb sequences, has also proven to be quite useful in predicting perturbations, including CRISPRi-mediated repression^65,71^. Further, several neural network models trained on human genomic sequences have shown that they can predict how variants impact gene expression to some degree when considered in aggregate, but are generally unreliable for individual variants^65,71,72^, and more reliable for promoter than for enhancer variants^71^. While such models are quite useful for many applications in their current state, they remain quite far from the models we require to reliably predict variant function. Given the large number of parameters needed to capture cis-regulation, building reliable quantitative models will require much more data.

### Why not just measure the genome more?

While we could continue functionally profiling the human genome, yielding ever more data and better models, we think this approach will soon reach a ceiling because of the limited length and sequence diversity of our genomes. Our genome consists of only ∼3 billion base pairs^73–75^. While ∼488 million base pairs appear to be cis-regulatory (open chromatin) in one or more cell types^76^, most cell types have ∼30 million cis-regulatory base pairs, little of which is specific to that cell type^76^. In contrast, if we were to query TF-TF interactions by instantiating one of every possible arrangement and offset for each TF-TF pair, this would take ∼5.7 billion bp (**Fig. 1B**)). Even if we had a genome comprised entirely of these TF-TF interaction query sequences, we still could not learn the interactions because multiple independent examples are required to distinguish the interaction-specific sequence features from those that are merely present. Accordingly, our genomes are simply too short to learn the many parameters describing cis-regulation.

In machine learning, it is important for the data to be independent and identically distributed (**i.i.d**.), which enables robust generalization to new data. For DNA sequences, i.i.d. would mean that the DNA sequences in the dataset are unrelated (independent), and that the active sequence features are equally likely to occur in any given sequence (identically distributed). In reality, our genomes are far from i.i.d., and evolution has shaped them in ways that can seriously confound genome-trained models. Regions of homology exist throughout our genomes, originating from individual DNA sequences that have been duplicated to other genomic loci and subsequently diverged. Nowhere is this more obvious and problematic than with repetitive elements that are estimated to comprise 60% of our genome^77^. For instance, LINE elements are 6-8kb retrotransposons that, through ongoing retroposition, now occupy ∼21% of our genomes^74^. Indeed, our genomes are so repetitive that the vast majority of pairs of 200 kb human genomic sequences from distinct chromosomes share thousands of 20 bp sequences (20-mers), with a median of >22,000 shared 20-mers representing a minimum of 11% overlap (if all 20-mers overlap maximally; **Fig. 2A-C**), despite the fact that 20 bp would be sufficient to uniquely specify each position in a non-repetitive sequence of 3 Gbp. Further, evolution favours clustering of related elements, with many loci containing multiple functionally related genes^78–80^, many genes having multiple enhancers with similar activity profiles^81–84^, and many enhancers having multiple binding sites for the same or functionally related TFs^85,86^. All this bias means that sequences sampled from the human genome are neither independent (due to homology) nor identically distributed (due selection favoring clustering of sequences with related functions). This will lead to the models trained on genomic data to mistake correlations for causation^71,87,88^ (**Fig. 2D,E**), resulting in unreliable predictions of regulatory element and variant function.

**Figure 2:**
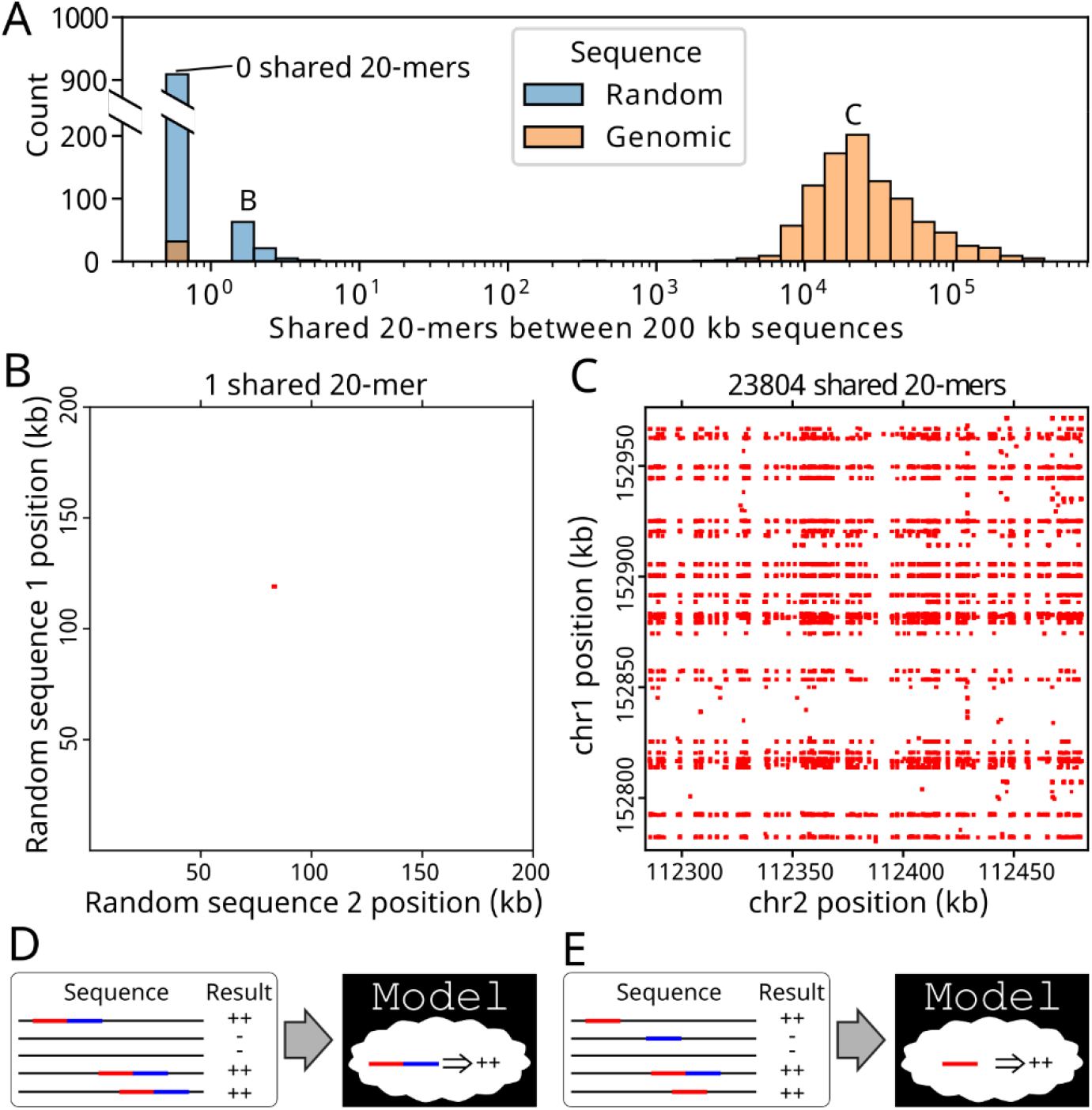
Training on the genome is confounded by cryptic homology and correlated but not causal features. (**A-C**) Cryptic homology makes data leakage nearly unavoidable with genomic DNA. (**A**) Histogram of the numbers of sequence pairs (*y*-axis) with each number of shared 20-mer matches (*x*-axis) between pairs of random (blue) and pairs of genomic (orange) 200 kb sequences (1000 pairs, each). (**B**,**C**) Examples of 20-mer matches (red boxes) between pairs of sequences (*x*- and *y*-axes) for (**B**) random sequences and (**C**) genomic sequences. (**D**,**E**) While confounded input sequence features can lead to models mistaking correlation for causation (**D**), causality is correctly inferred from random sequences (**E**).

As a consequence of the many biases and dependencies of the human genome, determining the true predictive power of models trained on genomic sequences is a major challenge. Model training is an ongoing race between fitting, where the model is learning things that are generally true about the system, and overfitting, where the model is learning spurious relationships that happen to be present in the training data. “Validation” data are used to determine when the rate of overfitting has surpassed the rate of fitting (when validation performance stops improving), at which point we should stop model training. Finally, “test” data are used to gauge the model’s true performance. In general, training, validation, and test data should be separate and distinct to ensure that overfitting and model performance estimates are accurate. Overfitting is surprisingly difficult to detect and avoid for genome-trained models^88–91^. Even initializing a model with TF motifs that were derived from the entire genome (which includes the test data) can lead to undetected overfitting ^92,93^. A major source of overfitting for genome-trained models is the abundant homology across the genome (**Fig. 2**), which makes it difficult - if not impossible - to avoid shared sequences in the training, validation, and testing datasets (termed “data leakage”), even when holding out entire chromosomes. Shared sequences in training and validation data will mask overfitting since the peculiarities the model has memorized from the training data are also present in the validation data, increasing the model’s performance on the validation data despite being overfit^89,90^. If these sequences are also shared with the test data, the overfitting will go completely undetected since the overfitting will improve test data performance too. For instance, rather than learning the biochemical mechanisms underlying LINE element activity, the model could simply associate LINE element activity to the most conserved sequence feature of the element. Because the model’s understanding of reality is flawed, it would then predict activity incorrectly for sequences that use similar biochemical features or have similar sequences to LINEs. Worse, this would go undetected if the training and test data both contained LINE elements, since it would predict their activity correctly, but for the wrong reasons (**Fig. 2D**). While one could mask repeat elements in an attempt to prevent data leakage, deciding which elements to mask is somewhat arbitrary since there is no level of divergence at which sequences sharing a common origin cease to be homologous. Further, there are many homologous sequences that are not derived from repeats (e.g. gene duplication, microsatellite repeats). Finally, the repeat elements are often active^94,95^, and so masking them would also necessarily confound model training and prediction. Even if one could avoid this data leakage, we will always need at least two instances of every genome-trained model, so that the predictions for a sequence come from a model that has not been trained on that sequence. Finally, the propensity for functionally related elements to cluster can also result in models having seemingly correct predictions but for the wrong reasons. For instance, if all enhancers within an enhancer cluster encode similar activities^81–84^ the model could ignore all but one of these elements, failing to accurately model the locus but correctly capturing the expression program. Consequently, it is incredibly difficult to get an accurate view of genome-trained model performance when evaluating them on genomic sequences (where we care about performance most).

There are several ways of augmenting the human genome sequence with other genome-derived sequences to improve models, but none will resolve the issues we identified above. For instance, training models on orthologous genomes (e.g. other mammals) may enable us to learn some of the rules of gene regulation, but issues of data leakage between orthologous genomes and our own remain even if direct orthologs are avoided. Further, some of the nuances of the cis-regulatory code are likely to be specific to humans (e.g. retrotransposon repressors, TF-TF interactions), and so would be learned incorrectly using this approach. Alternatively, we can train models with data designed to capture variant effects, for instance, using data from chromatin or gene expression profiling across people of diverse genetics and then learning genotype-aware models, or using measured expression effects of variants from Massive Parallel Reporter Assays (**MPRAs**)^96–99^. While training models with examples of variant effects will undoubtedly improve their performance, the cryptic homology causing data leakage remains problematic.

Ultimately, we want our models to learn all the causal relationships between sequence and expression while ignoring the spurious relationships that will lead us astray. Commonplace cis-regulatory mechanisms (e.g. billboard-like) will be learnable from genomic sequence because they are used abundantly throughout the genome in a variety of contexts. However, the genome’s abundant homology places a limit on the rarity of mechanisms that can be learned from its study; models will learn to associate the abundant spurious relationships before the rarer causal relationships (e.g. TF cooperativity) because they have no way of distinguishing between the two. Importantly, because of the issues described, determining the degree of overfitting of genome-trained models will remain a major challenge and we may never know exactly how much we can trust their predictions.

### Synthetic DNA as a potential solution

An alternative to training gene regulation models on genomic sequences is to train them on designed DNA sequences. Sequences can be designed in a variety of ways, including to test hypotheses about cis-regulation^25,32,100–102^ or its evolution^14^, or to achieve particular expression objectives^14,61,103^. The flexibility in design choices make designed sequences very powerful, but their comparatively high cost (∼$0.16 per 300 nt sequence as of 2023) limits the scale at which they can be applied. Learning cis-regulatory models from designed sequences can be challenging for several reasons. Firstly, the designed sequences may contain functional elements that were not intentionally added (e.g. cryptic binding sites^25^), and these must be accounted for. Secondly, it is common to use only a few different sequences to represent a TF’s binding site^32^, but models trained on such data will likely have difficulty predicting the effects of TF binding sites that do not resemble those designed. Finally, data leakage is easy with designed sequences resulting from shared features added by the researcher (e.g. the exact same TF binding site sequence is included in both training and test sets, or the designed sequences incorporate elements derived from the genome).

An alternative to designed sequences is to use random sequences, which has numerous advantages. The cost of the random DNA is negligible because it can be synthesized in a pool by a random process (e.g. ∼2,000,000,000,000,000 unique ∼120 bp DNA sequences for ∼$100). Since independently sampled random sequences share no more sequence identity than expected by chance, they are trivially divided into independent training, validation, and test datasets without fear of data leakage (although one must take care to account for errors in PCR and sequencing if these data were generated in the same experiment). Testing random sequences has proven to be an effective strategy for isolating functional nucleic acids^104^ and proteins^105^, learning 5’UTR ^106,107^, splicing^108,109^, and polyadenylation logic^110^. Importantly, random sequences have developmental enhancer activity^111^, and we recently showed that random DNA provides ideal data for deciphering cis-regulation^14,25,31^. For instance, our (de Boer) recent proof-of-principle study in yeast showed that random DNA had diverse cis-regulatory activity, and this enabled us to design an experiment for simultaneous quantification of the expression levels encoded by each of >30 million random 80 bp sequences (∼2.4Gb of regulatory DNA; 184 yeast genome equivalents)^25^. The richness and abundance of these data gave us immense power to detect even relatively weak phenomena, including the position-dependence of TF binding on activity for all yeast TFs at base pair resolution^25^, which is simply not possible to learn by studying the yeast genome because of its limited sequence diversity. Inspired by this work, we (Taipale group) recently used a similar strategy in human cells, identifying transcriptionally active DNA sequences from a sequence space ∼100 times larger than the human genome. This demonstrated that random DNA has promoter and enhancer activity, enabling data generation at unprecedented scale for the creation of human cis-regulatory models^31^. The vast data one can generate with random approaches powers models to learn even rare features with modest effect sizes.

While learning cis-regulation from measurements of random DNA activity may seem counter-intuitive, the reason why this works is simple: there are many TFs (∼1,639 in humans^2^), and their binding sites occur frequently by chance^25,111,112^, enabling one to select for binding sites for individual TFs or pairs of TFs from random DNA^37,113,114^. While a random sequence having a binding site for a specific TF is unlikely, it is almost certain to have binding sites for some TF, leading to high expression diversity among random sequences^25^, and distinct expression programs across cell types^111,115^. This propensity for random sequences to frequently be active is also consistent with the soft regulatory syntax of the billboard model^23^. The comparative ease at generating regulatory sequences from random DNA is also supported by evolutionary study: if it were difficult to generate TF binding sites or active regulatory sequences by chance, we would expect that regulatory elements would evolve primarily through duplication and divergence, like the protein coding genes. However, this is not what is observed. Cis-regulatory elements turn over rapidly over evolutionary time^116^, often with little conservation of TF binding sites or enhancer sequences, despite the TFs and expression programs being conserved^117,118^. Thus, most regulatory sequences appear to have evolved from previously non-regulatory DNA^119–122^, demonstrating that the syntax by which TFs act must be relatively permissive such that functional elements can emerge by chance. Indeed, the proportion of random sequences with regulatory activity is similar to genomic sequences^25,31,115^. This rather counterintuitive finding makes sense from an evolutionary perspective since there is no advantage to increasing the specificity of a cis-regulatory element (let alone all elements) beyond what is required for its function, and the small marginal advantages to further specificity beyond this (if they exist at all) can be dominated by the effects of genetic drift^123,124^. Further, a cis-regulatory code that was so specific that it prevented regulatory innovation would prevent organisms from adapting to new regulatory optima (e.g. a new environment). While highly optimized regulatory elements like the IFN-B “enhanceosome”^125,126^ exist, they represent the exception, and could not have evolved de novo in their current form; evolution must proceed one mutation at a time (**Fig. 3A**). Each mutation in the evolution of a complex cis-regulatory element must make use of the same biochemistry used by newly-evolved enhancers and active random sequences^127^ (**Fig. 3**). Reporter assays of random DNA exploit the propensity for regulatory sequences to occur by chance to enable quantification of cis-regulatory sequences at unprecedented scales^25,31^.

**Figure 3:**
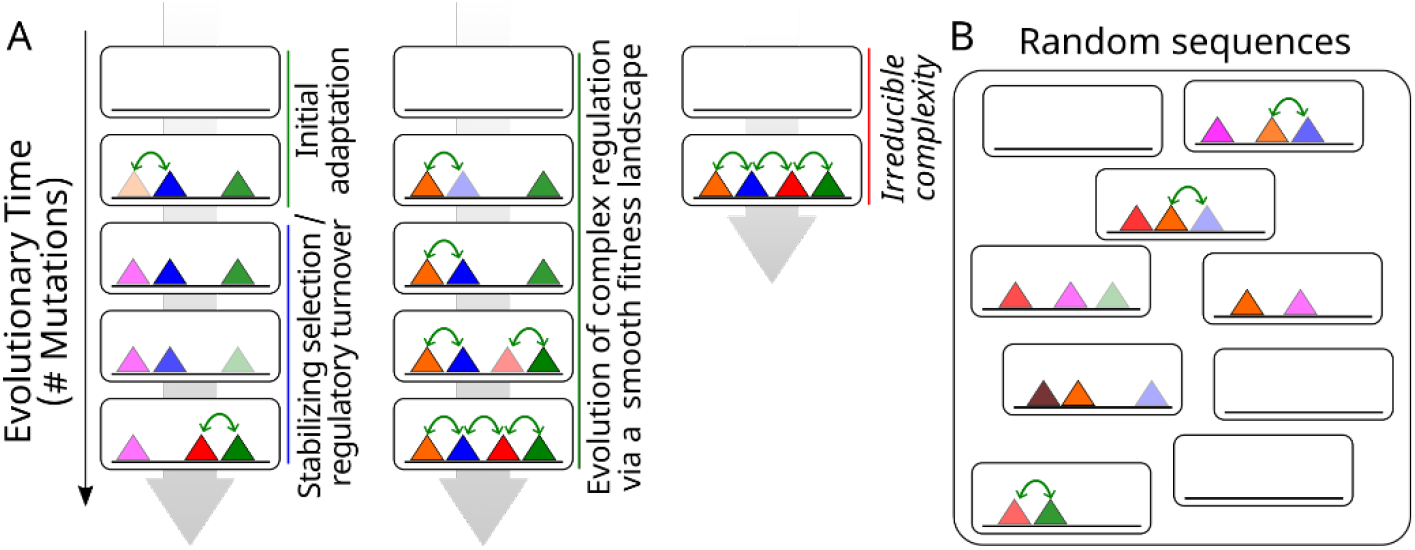
Evolved and random regulatory elements use the same biochemical rules. (**A**) Most regulatory elements appear to evolve de novo from previously non-regulatory DNA (left). Random mutations in a previously non-regulatory sequence produce an advantageous initial function. Once established, there is often little advantage to increasing in complexity, and the mechanisms used by the regulatory element can drift so long as its activity is maintained^14,117^. In some cases, complexity can evolve where precise control of a regulatory element is advantageous (middle), but it too evolves via the same random mutations and biochemical rules. Meanwhile, it is highly improbable for such a complex sequence to evolve from scratch (right). (**B**) Random sequences are biochemically active via the same mechanisms as evolved sequences.

A second counter-intuitive aspect of using random DNA to learn the cis-regulatory code is that the code can be broken despite the experimentally measured random DNA representing a vanishingly small fraction of the possible sequence space. In science, we have been trained to systematically test all possibilities to determine the impact of each variable in the system. Random DNA sequences achieve this isolation of variables precisely because the sequence is random. Features, such as TF binding sites, are uncorrelated with their flanking sequences, but are correlated with the regulatory effect of that TF’s binding. The only requirement to learn these causal relationships from random DNA is that we have sufficient instances of them in the training data to attribute the observed activities to the functional elements that generated the activity.

Regardless of training data source, the ultimate test of models is their ability to predict genomic activity. While still at the early stages, our previous work has already illustrated that training cis-regulatory models on random DNA sequence activities can produce robust models that predict the activities of genomic sequences (and variants thereof) from sequence alone^14,25,31^. These models can surpass the performance of genome-trained models by having access to orders of magnitude more data than the genome can provide. Importantly, models that can predict genomic activity but have never seen genomic sequences before cannot be cheating via undetected overfitting, and so we will be able to gauge their performance and trust their predictions much more than genome-trained models.

### Current and needed technologies to facilitate large-scale synthetic regulatory DNA assays

For both random and designed sequences, recent advances in DNA synthesis and reporter assays have enabled us to quantify the expression of non-genomic sequences at scales surpassing the genome. While we have had success with random sequences where each base is randomly sampled from the four nucleotides to achieve an overall base content similar to the genome^25,31^, random sequences can also be biased in certain directions, if needed. For instance, the base content (e.g. dinucleotide frequencies) can be adjusted or certain positions can be held fixed to increase the frequency of specific TF binding sites or adjust the proportion of active sequences. Designed sequences are much more expensive and limited in scale, but are extremely powerful because they enable researchers to test whatever sequences they like. For instance, one can use computer models to generate sequences designed to maximally improve the model (**Active Learning**). This could include sequences that are predicted to have extreme biochemical activity that are rare in both random and genomic sequences. With measurements from these extreme sequences, we can create superior models that have never trained on genomic sequences. While random sequences have the advantage that they can easily and cheaply explore the sequence space, we anticipate that designed short DNA sequences will play an increasingly important role in training cis-regulatory models as DNA synthesis continues to get cheaper.

Technologies for measuring the activities of non-genomic sequences will also continue to evolve, enabling us to look at different aspects of the cis-regulatory code (**Fig. 4A**). Reporter assays, which aim to directly quantify the relationship between sequence and expression in a controlled system, have proven to be invaluable in understanding the cis-regulatory code due to their ease of application and relatively high throughput, and indeed most of the non-genomic cis-regulatory data that has been generated is via reporter assays. While we have several technologies for reporter assays, none are ideal. While STARR-seq^128,129^ can be applied to quantify tens of millions of sequences, it includes substantial measurement noise at this scale due to biases in quantifying the input DNA and output RNA for each sequence^31^. Further, transiently transfected DNA can initiate immune responses in the cells^130,131^, confounding the readout of the assay. Additionally, RNA-based reporter assays cannot capture several important aspects of transcriptional regulation that are thought to be important in disease, namely, transcriptional responses that happen at short time scales (e.g. transient TF activation^132,133^), and intercellular expression variance^134^. Stably integrated translated reporters^135^ and single cell MPRAs^136,137^ theoretically overcome many of these limitations, but have so far not been widely applied at scale. While reporter assays can produce rich data, it will also be useful to measure the biochemical intermediates between sequence and expression, for instance detecting TF binding using enzymatic footprinting^138^. In addition, combining reporter assays with perturbations of regulators^139–141^ will allow us to connect the identities of the regulators with their cognate functions in the models (**Fig. 4B**). Developing systems for testing more DNA in more ways and in more cell types will be an important area for future work.

**Figure 4:**
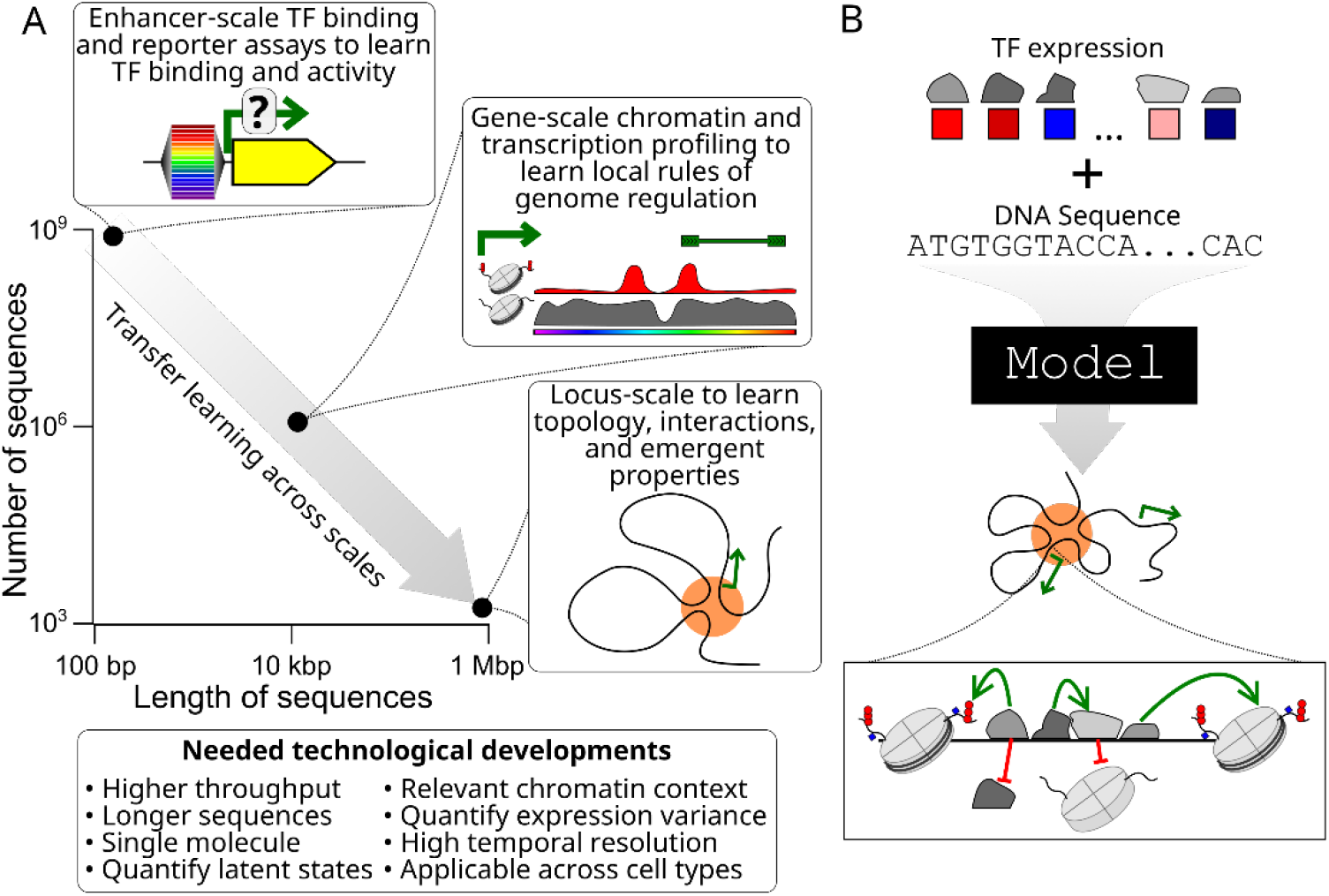
Current and future technologies to solve cis-regulation. (**A**) Different sequence lengths and scales are feasible for different measurement modalities. Using transfer learning, we can learn a single cis-regulatory model that incorporates regulatory mechanisms at different scales. (**B**) The resulting model would take as input a DNA sequence and TF expression (to capture cell types/states and associate TFs with cognate activities in the model) and would predict higher-level chromatin features based on a detailed understanding of cis-regulation.

While many of the low-level features of cis-regulation (e.g. TF binding) can likely be learned from short sequence reporter assays, ultimately, we must learn how regulatory elements all work together in the context of a chromosomal locus. Learning the higher-order regulatory mechanisms, including 3-D looping^49–51^, insulation^52^, competition^53–55^, and condensate formation^56,57^ will require assaying the regulatory activities of longer DNA sequences (**Fig. 4A**). However, both random and specific DNA synthesis of individual oligonucleotides are limited in length to ∼300 bp. Technologies for pooled gene synthesis (e.g. DropSynth^142,143^) enable construction of ∼1000s of sequences, each less than 1 kb, and so may be useful in testing small libraries of regulatory sequences, but this technology has not yet reached the scale where training data can be generated in sufficient quantities. It is now possible to synthesize ∼100 kb of designed DNA by assembly of smaller constructs^144–147^, and these have been used to study genome regulation^144,145^. While major efforts aim to make this more routine^148^, it is presently far too laborious and costly to produce the needed training data at a reasonable scale. Long random DNA sequences are predicted to be active in human cells^115^, suggesting that this strategy may work here as well. As before, specific elements could be spiked in to produce data more conducive to model training (e.g. higher frequencies of regulatory activity). Synthesis of random or partially random long DNA sequences could also be substantially less challenging than synthesis of specific sequences since most errors in synthesis, such as point mutations, insertions, and deletions are tolerable as they simply generate another random sequence. A second challenge is integrating the DNA into the genome, which will have to be made more efficient and accessible across cell types. A final challenge is developing the approaches for characterizing the activities of long non-genomic sequences. All of the same functional genomics approaches used on the genome could be used to study the activity of non-genomic DNA (e.g. RNA-seq, DNaseI-seq, ATAC-seq, ChIP-seq) by mapping reads to a hybrid genome containing both genomic and synthetic DNA. In order to understand the dynamics of the system, developing multiplexed readouts of chromatin activity, ideally at single molecule resolution, will become increasingly important^149,150^. In addition to providing the needed data with which to train our genomic models, the scaled assay of long random sequences would also help to define a genomic null hypothesis describing how frequently we expect to see regulatory activity by chance^115,151^.

### Moving forwards

The limitless supply of non-genomic sequence and its frequent activity make testing non-genomic sequences in high throughput a viable path for solving cis-regulation. Using short-sequence genome-integrated reporters and related assays of both random and designed sequences in extremely high throughput (>10^9^ separate elements), we will be able to train cis-regulatory models that understand the biochemistry that underlies TF binding and gene expression in the context of chromatin. We can then use transfer learning, where parameters learned in one system are used to prime models being trained to capture another, to adapt what we have learned on shorter sequences to longer DNA sequences with more complex regulation (which cannot yet be assayed in as high throughput) and inaccessible cell types (e.g. using scATAC-seq data; **Fig. 4**). Ultimately, we want models for every cell type in the human body. Building the genetic and experimental tools we need to generate non-genomic training data in pluripotent cells provides a straightforward way to enable quantification of non-genomic sequence activities across cell types and throughout development. Accordingly, it will be important to continue to develop systems for cellular differentiation and reprogramming^152,153^. Since assaying non-genomic DNA at large scale across development remains a monumental task, a coordinated effort from the genomics community is needed.

Solving the sequence-to-expression problem will also require continued development of computational models. In particular, approaches to integrate data across scales, approaches, and cell types, including by transfer learning or multimodal prediction, would enable models to integrate evidence from diverse types of input data. For instance, integrating data from in vitro TF binding (e.g. SELEX), reporter assays, and chromatin measurements from long sequences would enable the models to learn the parameters shared across them more efficiently. In the short-term, transfer learning from non-genomic sequence models to genomic sequence models will likely improve their performance since the spurious genomic associations are absent in the non-genomic sequence^156^. Once we have sufficient data, we can use transfer learning from short non-genomic sequences to long non-genomic sequences to have fully genome-independent cis-regulatory models. Further development of protein-structure/affinity prediction models that can predict protein-protein^154^ and protein-DNA complexes ^155^ that form on DNA in a sequence-dependent manner will also enable us to encode priors for some of the relevant cis-regulatory parameters, increasing the predictive power of the cis-regulatory models. Finally, having standardized datasets and modeling competitions (e.g. DREAM Challenges) will facilitate the continued improvement of model efficiency and accuracy.

While the human genome is too small and too repetitive to learn cis-regulation in the necessary detail, continuing to characterize it will be important for generating the datasets required to validate cis-regulatory models. Establishing Gold Standard genomic datasets will enable the field to identify model deficiencies, highlight areas for improvement, motivate experiments designed to complement the existing data, and identify new biological phenomena. Measurements of cis-regulatory variant effects are particularly useful since predicting their effects is a major motivation for model improvement. In the near term, we should continue to develop models trained on genomic data to further our understanding of genome regulation in health and disease, keeping in mind their limitations and trying our best to avoid data leakage. As more and more non-genomic data are generated, we can move the genomic data to the test set, where it will help us determine when our knowledge of cis-regulation is complete. When we can predict the activity of genomic sequences and mutations to them to the degree that is explainable given the limitations of the assays using a model that has never seen genomic sequence before, we will know that cis-regulation is solved.

## Acknowledgements

We would like to thank Brian Cleary, Abdul Muntakim Rafi, Omar Tariq, Sara Mostafavi, Xinming Tu, Alex Sasse, Bas van Steensel, Benjamin Lehner, Anshul Kundaje, Matthew Weirauch, and Gökçen Eraslan for helpful discussions. We apologise to all our colleagues whose work we could not cite due to the limit on the number of references.

## Methods

### Shared 20-mer analysis

Random sequences were generated with bases sampled randomly using the human genomic average (41% G+C). Chromosomal sequences were sampled from random genomic locations (GRCh38) ensuring that any pair being compared were on different chromosomes, and that there were no Ns in the sampled DNA sequences. 1000 pairs of random sequences and 1000 pairs of chromosomal sequences were generated, with each individual sequence 200 kb long. For each pair, the numbers and locations of common 20-mers were identified, considering 20-mers shared between the first sequence and either the forward or reverse complemented second sequence. A length of 20 was chosen because a de Bruijn DNA sequence, which includes all possible k-mers without repetition, for k=20 is substantially larger than the size of our genomes (10^12^ bp).

